# Extralimital Terrestrials: A reassessment of range limits in Alaska’s land mammals

**DOI:** 10.1101/2023.11.01.565160

**Authors:** A. P. Baltensperger, H. C. Lanier, L. E. Olson

## Abstract

Understanding and mitigating the effects of anthropogenic climate change and development on species requires the ability to track distributional changes over time. This is particularly true for species occupying high-latitude regions, which are experiencing a greater magnitude of climate change than the rest of the world. In North America, ranges of many mammals reach their northernmost extent in Alaska, positioning this region at the leading edge of climate-induced distribution change. Over a decade has elapsed since the publication of the last spatial assessments of terrestrial mammals in the state. We compared public occurrence records against commonly referenced range maps to evaluate potential extralimital records and develop repeatable baseline range maps. We compared occurrence records from the Global Biodiversity Information Facility for 64 terrestrial mammals native to mainland Alaska against a variety of range estimates (International Union for the Conservation of Nature, Alaska Gap Analysis Project, and the published literature). We mapped extralimital records and calculated proportions of occurrences encompassed by range extents. We also evaluated extralimital records against published species models, highlighted extralimital observations on U.S. Department of Interior lands, and report on two species of bats new to Alaska since 2014. Range comparisons identified 6,848 extralimital records belonging to 39 of 112 (34.8%) Alaskan species. On average, the Alaska Gap Analysis Project encompassed 95.5% of occurrence records and ranges were deemed accurate (> 90.0% correct) for 31 of 37 species, but overestimated extents for 13 species. International Union for Conservation of Nature range maps encompassed 68.1% of occurrence records and were > 90% accurate for 17 of 39 species. Results are either the product of improved sampling and digitization or represent actual geographic range expansions. Here we provide new data-driven range maps, update standards for the archival of museum-quality locational records and offer recommendations for mapping range changes for monitoring and conservation.

## INTRODUCTION

Quantifying the degree to which species ranges have shifted over time is essential for understanding organismal responses to anthropogenic climate change. Although these efforts are key for biodiversity conservation [1] documenting, quantifying, and predicting such shifts is challenging, particularly given the rapid pace of global change. For example, the effects of climate change at high latitudes are especially pronounced [2] with mean annual and seasonal temperatures increases of 2-4°C over the past 20 years resulting in thawing permafrost, changing precipitation patterns, advancing shrub- and tree lines, and shorter fire-return intervals [3]. The disproportionate effects of climate change at high latitudes are already causing a northward shift of the boreal forest at the expense of Arctic tundra, reallocating available habitat for Arctic- and boreal-adapted species [4–6]. Although many species in these biomes are expected to maintain habitat associations and move in accordance with shifting landcover [7–10], sufficient spatial data are lacking for many species [11] making shifts in species’ ranges challenging to establish. Understanding the impact of these habitat shifts on community assemblages requires accurate approaches for mapping and assessing species ranges.

Tracking species and community range shifts requires knowledge of prior species distributions and assessment of putative extralimital occurrences. Wildlife ranges are inherently dynamic, even during periods of relative stasis [12,13], so distinguishing true distributional shifts from newly collected information at poorly characterized range margins can be challenging. Thus, the identification of new extralimital occurrences is highly dependent on the range to which they are compared. One approach for developing a background range is to use species distribution models (SDMs) or ecological niche models (ENMs), which can incorporate environmental variables in often discontinuous predictions across landscapes [14–16]. However, SDMs often lack repeatability, methods can vary considerably among studies and over time, and models may predict suitable but as-yet unoccupied marginal habitats [17,18]. In contrast, the use of occurrences to delineate range extents (i.e., classic range maps) and identify extralimital records (i.e., occurrences outside identified range extents) may be the most robust and repeatable approach. Admittedly, range maps are coarse and generalize by nature and therefore often overestimate real occupancy [19], but they represent consistent baseline summaries of existing knowledge, often from citable literature, against which new occurrences can be compared.

A further complication for identifying marginal shifts in species is that even verified occurrences indicate only that an individual of a species was found at a particular time and place and do not necessarily suggest regular habitation by that species [20]. Furthermore, due to potential sampling biases or low marginal abundances, extralimital records could reflect improvements in sampling and knowledge of occurrences near range boundaries. Nevertheless, without developing accurate baseline range extents, monitoring species change across space will remain an unmet challenge. Ultimately, regular, geographically representative sampling that enhances comprehensive occurrence datasets is the best scientific means for monitoring changing range extents in the context of rapid climate change [1].

The ranges of numerous North American mammals reach their northern- and westernmost extents in Alaska, and so range limits in the state also take on continental and, for some, global significance [21]. As we monitor changing species distributions and community assemblages, quantifying the dynamics of species near the leading edges of expanding ranges is paramount, as this space also represents the receding edge for several Arctic endemics whose ranges are likely contracting [4,22–24]). Climate change in Alaska is predicted to cause the geographic ranges of species to shift northward, upward in elevation, and towards the coast [4,23,25]), yet the status of such predicted species-specific movements remains largely unknown. Limited research has examined the extent to which terrestrial mammalian range or distribution shifts have actually occurred for select species over historical timescales in Alaska, including the expansion of American Marten (*Martes americana* [Turton, 1806]) across the Kenai Peninsula [26], northward shifts in the Alaskan Hare’s (*Lepus othus* [Merriam, 1900]) distribution in western Alaska [27], and the expansion of Snowshoe Hares (*L. americanus* [Erxleben, 1777]) [28], Moose (*Alces alces* [Linnaeus, 1758]) [29,30], and North American Beavers across the Alaskan Arctic [31].

Several research efforts conducted in the 2000s endeavored to collate and map occurrences of mammals in Alaska [21] and describe ranges of terrestrial vertebrates and other species (e.g., [32,33]). MacDonald and Cook [21] compiled georeferenced museum specimens for recent marine and terrestrial mammals in Alaska and mapped occurrences and marginal records across the state. This marked a notable milestone in Alaskan mammalogy, as no such effort had been attempted previously, and the resulting maps remain the standard against which new occurrences are interpreted. Between 2009 and 2014, the Alaska Gap Analysis Project (<u>AKGAP</u>) conducted a spatial review of distributions and range limits for all terrestrial vertebrates in Alaska using occurrence records and three geospatial analyses [33]. The International Union for the Conservation of Nature (<u>IUCN</u>) has assessed conservation threats since 1964 for more than 150,300 species worldwide and also serves range maps at a global scale used to inform the conservation status of vulnerable species on IUCN’s Red List [32].

Since 2009, advances in digital data collection and the rise of online museum and citizen-science initiatives such as <u>iDigBio</u> and <u>iNaturalist</u>, respectively, have enhanced field detection of species and significantly augmented the publicly available, georeferenced datasets from museums (e.g., <u>Arctos</u>, <u>VertNet</u>). Citizen-science applications, like iNaturalist, allow observers to easily contribute georeferenced, photo-verified records of species from a variety of personal devices (complete with coordinates and metadata). If a plurality of users agrees with the species identification, photo-observations may be deemed “research grade” and uploaded to the Global Biodiversity Information Facility (<u>GBIF</u>). GBIF is the world’s largest and most comprehensive digital biodiversity repository (> 2.3 billion records) and it regularly harvests and collates records from iNaturalist and from dynamically curated, specimen-based databases like Arctos, VertNet, <u>BISON</u> (Biodiversity Information Serving Our Nation), and iDigBio, which have been steadily digitizing analog museum records.

The wealth of new, georeferenced, and digitized specimens, as well as the growth in research-grade citizen science data, provides a timely opportunity to re-examine occurrence datasets of Alaskan mammals in relation to range estimates over the past decade. Here we provide an update to the geographic status of northern terrestrial mammals with records beyond previously defined range extents. We review locational records [21,34] and range maps (IUCN and AKGAP) for 64 Alaskan mammals. We place new extralimital records in the context of federal lands, highlight species newly documented in Alaska, correct dubious records, evaluate the accuracy of commonly used range maps, contribute new, current, repeatable range maps, and review predictions made by previously developed future distribution models. We also reassess the standards of identification, archival, and public accessibility of georeferenced occurrence records. We undertook these analyses with the goal of improving the quality of source data underpinning range maps and predictive models, so that occurrence record sets and current range maps can provide conservation planners and wildlife managers with improved means for monitoring changing terrestrial wildlife patterns in Alaska.

## MATERIALS AND METHODS

To obtain the most comprehensive set of publicly available mammal occurrence records, we searched GBIF for all georeferenced (coordinate precision < 5 km) records (i.e., whole specimen, tissue sample, machine observation [remote detection], human observation) of terrestrial mammals known or suspected to occur in Alaska, USA [21,34]. Records were not filtered to exclude duplicate observations of a species at the same location. We followed the taxonomy used by the American Society of Mammalogists Mammal Diversity Database (ASM; mammaldiversity.org/explore) and evaluated 64 native, terrestrial, non-island-endemic, mammal species (including bats) thought to occur naturally (i.e., not introduced) in Alaska (Tables 1, S1 Table; [21,33]. Baseline (2009) range polygons were constructed in ARCGIS 10.5 (ESRI, Inc., Redlands, CA) by digitizing marginal records for each species using coordinates provided by MacDonald and Cook [21] and then constructing minimum convex polygons (MCP) from the data.

In ARCGIS, we intersected species-specific GBIF record sets with their corresponding MCP to identify and tally extralimital records (those occurring at least 5 km outside baseline range margins; [21]) and used these species for further analysis (Table 1). We flagged records with suspected georeferencing or taxonomic errors as dubious and in need of further clarification, correction, or omission, and institutions of origin were consulted accordingly. Records that were confirmed as accurate and those that were corrected by museum professional met our criteria as “verified” and were included in final species occurrence datasets, whereas those remaining with implausible dates or locations were deemed unreliable and excluded from further analysis.

**Table 1.**
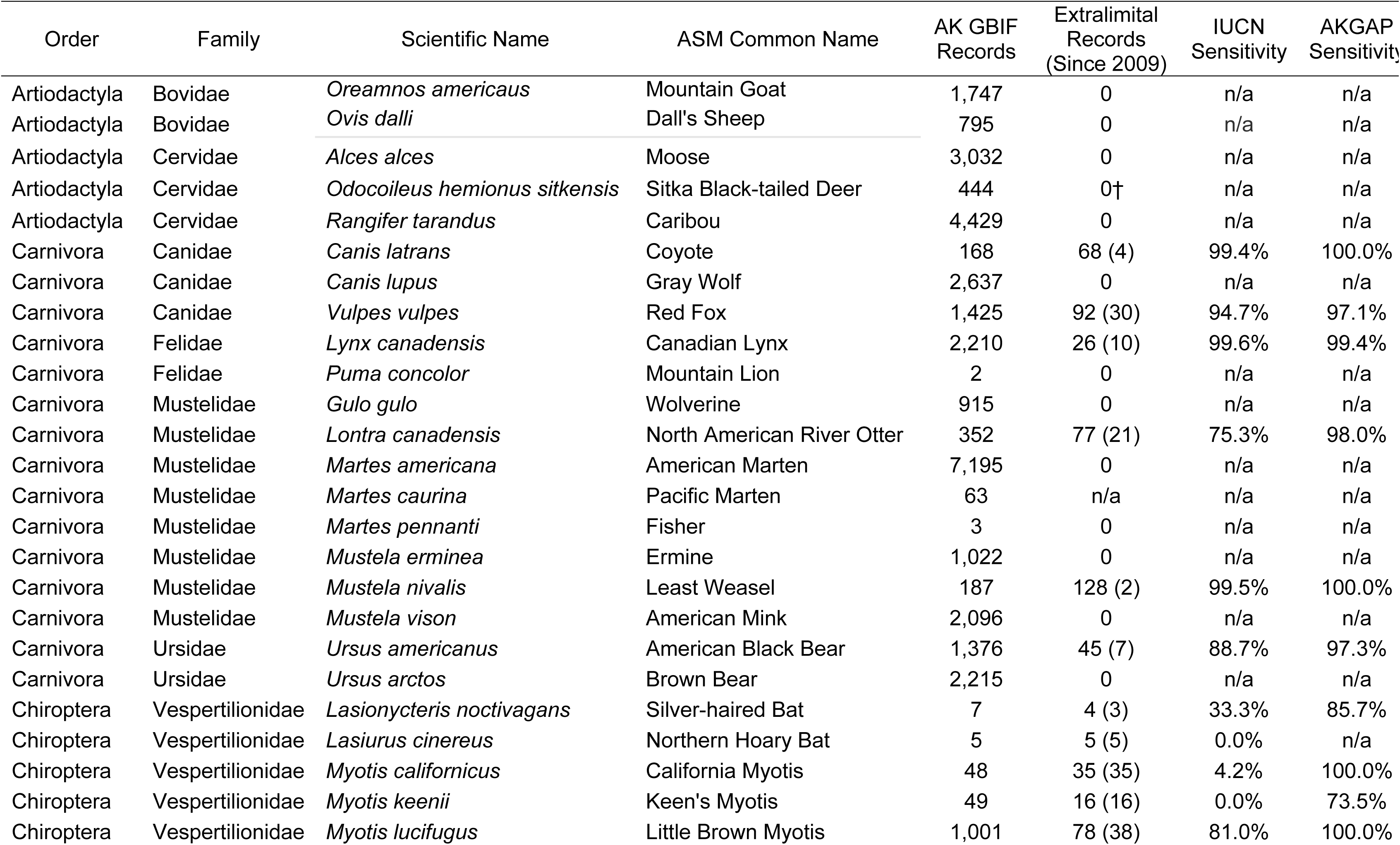
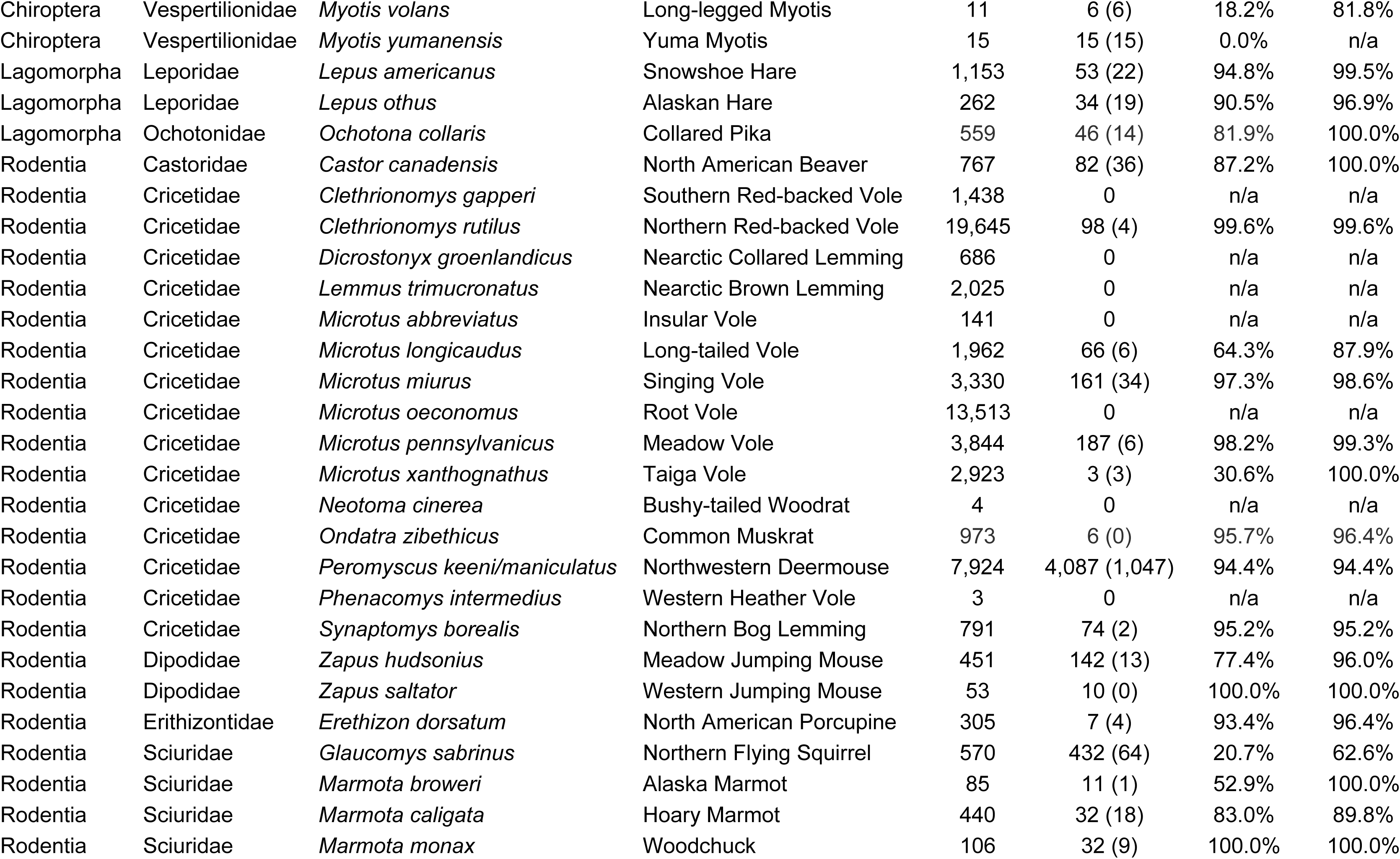
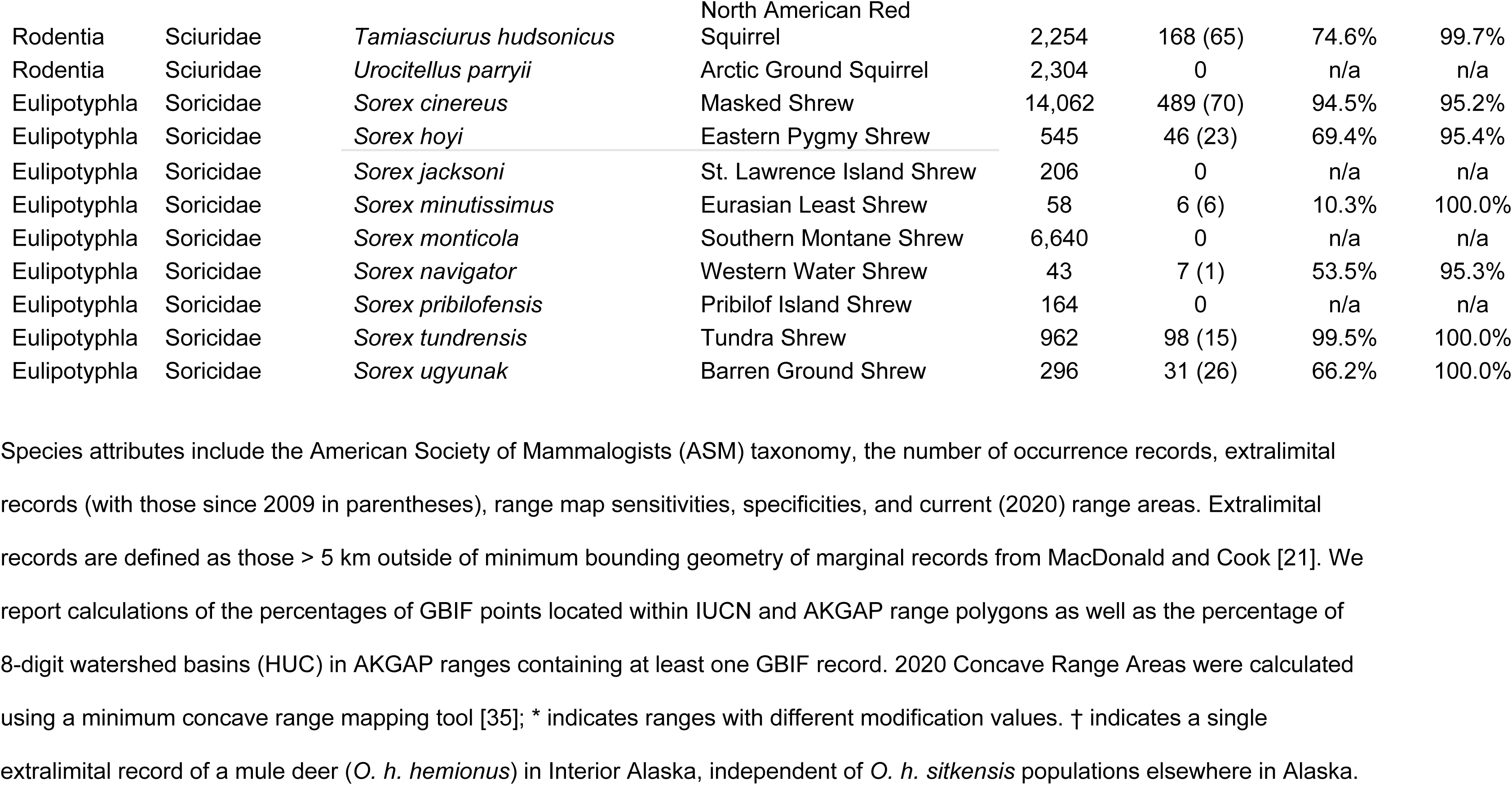
List of terrestrial mammal species in Alaska.

Species attributes include the American Society of Mammalogists (ASM) taxonomy, the number of occurrence records, extralimital records (with those since 2009 in parentheses), range map sensitivities, specificities, and current (2020) range areas. Extralimital records are defined as those > 5 km outside of minimum bounding geometry of marginal records from MacDonald and Cook [21]. We report calculations of the percentages of GBIF points located within IUCN and AKGAP range polygons as well as the percentage of 8-digit watershed basins (HUC) in AKGAP ranges containing at least one GBIF record. 2020 Concave Range Areas were calculated using a minimum concave range mapping tool [35]; * indicates ranges with different modification values. † indicates a single extralimital record of a mule deer (*O. h. hemionus*) in Interior Alaska, independent of *O. h. sitkensis* populations elsewhere in Alaska.

We focused subsequent analyses on those species with extralimital records (as defined in the previous paragraph; Table 1). For each species we intersected verified occurrences in ARCGIS with corresponding IUCN and AKGAP range maps [32,33] to determine the degree to which these maps corresponded to occurrence datasets. To evaluate the specificity of AKGAP ranges, we also calculated the percentage of 8-digit hydrologic units (HUCs) within the ranges of species containing at least one occurrence record. All intersections used a spatial tolerance of 5 km. We also identified species with new extralimital records that occurred in units managed by the U.S. National Park Service (including National Parks [NP], National Preserves [which we do not abbreviate here for the purposes of disambiguation with National Parks], National Historic Parks (NHP), and National Parks and Preserves [NP&P]) and the U.S. Fish and Wildlife Service (including National Wildlife Refuges [NWRs]; Fig 1), two federal agencies whose holdings are uniquely situated to monitor range shifts on federal lands in Alaska.

**Fig 1.**
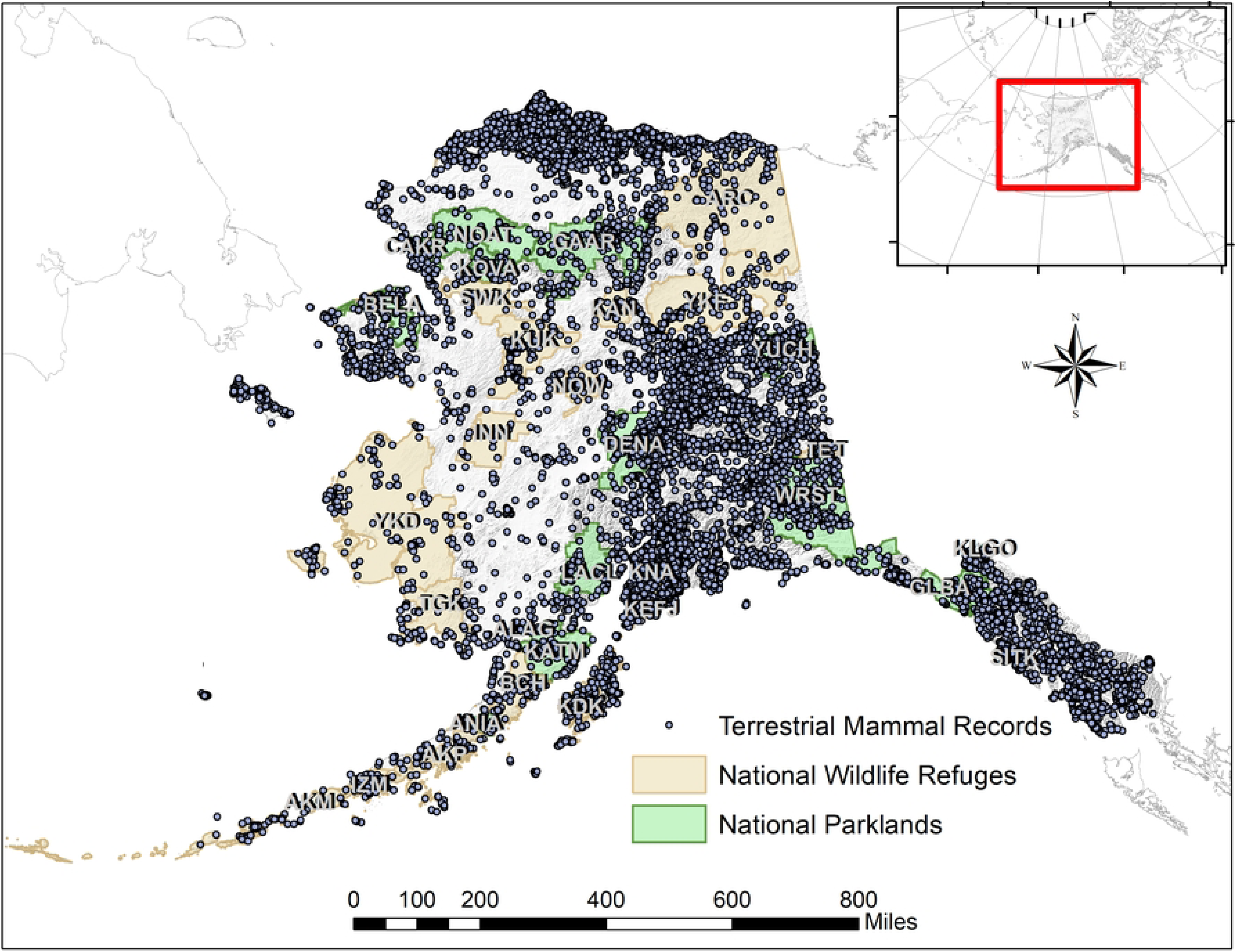
Map of Alaska showing all records of native, terrestrial mammals downloaded from the Global Biodiversity Information Facility in relation to U.S. National Park Service and U.S. Fish and Wildlife Service managed land units and their standard abbreviations (See Table 2).

**Table 2.**
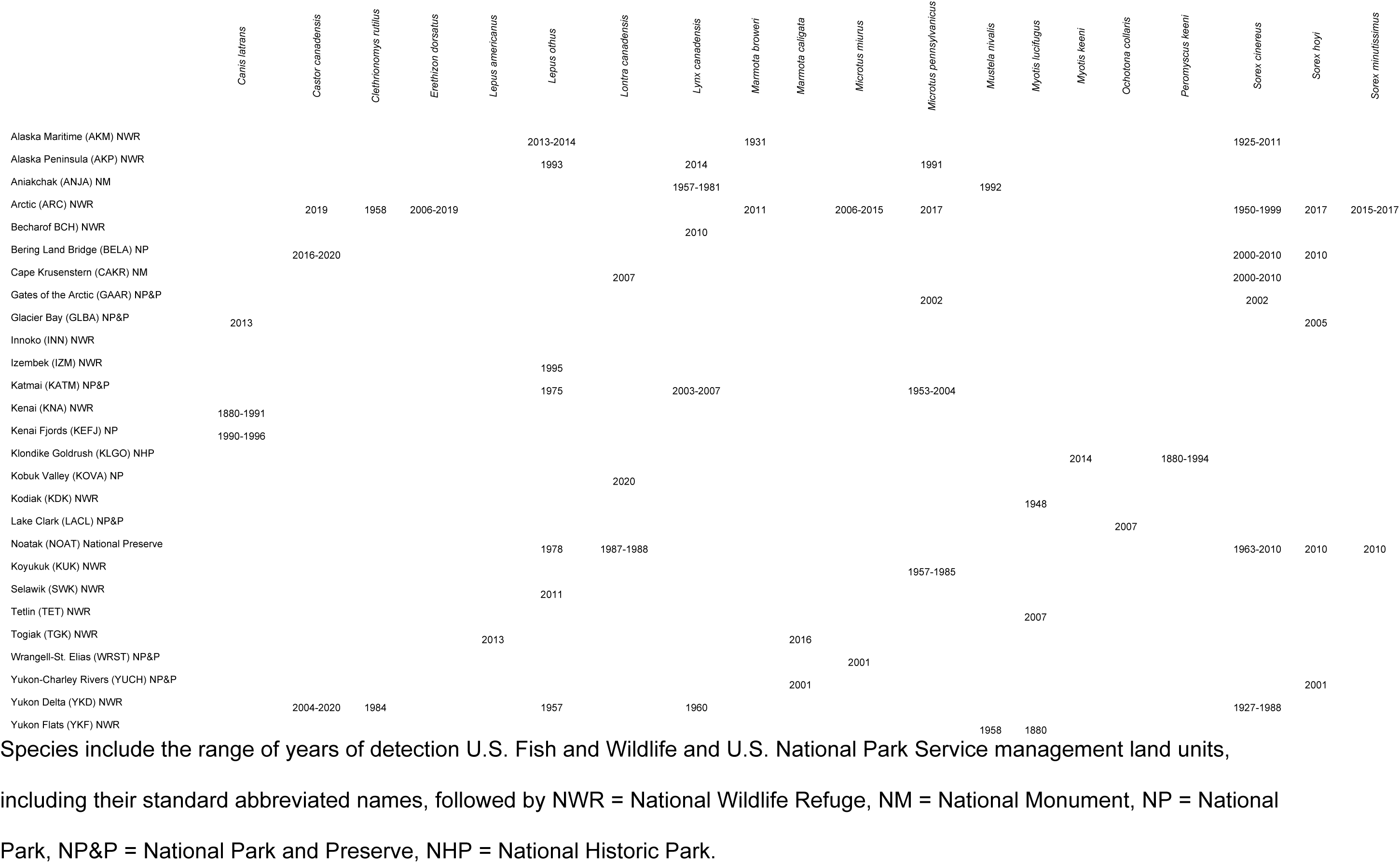
Terrestrial mammalian species with extralimital records.

We subsequently developed simple range models for the 39 species with extralimital records, using our updated geodatabase of occurrence records. We used the Python tool, Concave Hull from Occurrence Points [35,36], to define concave polygons, representing current, inferred range extents for each species. Tool performance and range geometry is based on a minimum distance setting and a scale-dependent multiplicative factor that adjusts the aggregation distance to produce repeatable maps that do not over generalize [36,37]. We used the default minimum search distance of 1,000 m, a buffer distance of 5 km, and a modification value of 12 for all species except Singing Voles (*Microtus abbreviatus* [G. S. Miller, 1899]) and Long-tailed Voles (*Microtus longicaudus* [Merriam, 1888]), for which we set the modification value to 8 and 15, respectively, to account for their disjunct distributions.

## RESULTS

### Range Evaluations

From GBIF we harvested a total 121,943 records of 64 native, terrestrial, non-island-endemic mammals with occurrences in Alaska (Fig 1) and identified 39 species as having extralimital records beyond previous MCP range extents (Table 1) [21]. Comparing occurrences for those species to IUCN and AKGAP range maps, we found that, on average, IUCN range maps encompassed 68.1% of GBIF records, whereas 95.5% of GBIF records were correctly included in AKGAP range extents (Table 1). AKGAP ranges exceeded 90.0% accuracy (correctly included presences / all occurrences) for 31 of 37 species (83.7%), whereas IUCN range maps reached 90% accuracy for just 17 of 39 species (43.6%) with available maps (Table 1). However, AKGAP ranges overestimated range extents for most species by including numerous 8-digit HUCs without occurrences. Among reviewed AKGAP range maps, an average of 57.3% of HUCs contained GBIF records while HUC occupancy exceeded 50.0% for 21 of 37 species (56.8%; Table 1).

### Extralimital Records

We identified 39 species as having ≥1 extralimital records since 2009 (Table 1, Fig 2). From these we identified 6,853 extralimital records (including multiple specimens from the same locations) belonging to species from five orders (Eulipotyphla, Rodentia, Lagomorpha, Chiroptera, and Carnivora; Tables 1-2, Fig 2, S2 Table). Among these, 1,702 extralimital detections occurred in 2009 or later. Of the extralimital records, 3,932 belonged to a single species, the Northwestern Deermouse (*Peromyscus keeni* [Rhoads, 1894]), whose taxonomy is entangled with that of the Eastern Deermouse (*P. maniculatus* [J. A. Wagner, 1845]). We flagged an additional 310 records (4.5%) that appeared to be extralimital, but which were incorrectly georeferenced or misidentified taxonomically as dubious. Through communication with museums, we resolved georeferencing errors for these specimens. Below, we describe the extralimital dataset by taxonomic order with a focus on how recent extralimital (≥ 2009) records compare to prior range limits (Table 1, Fig 2) and how they are distributed across lands managed by the U.S. National Park Service and U.S. National Wildlife Service (Table 2, Fig 1). Additional information and datasets are available in the Supplementary Files (S1-S2 Tables and at Open Science Framework (https://osf.io/h6u5n).

**Fig 2.**
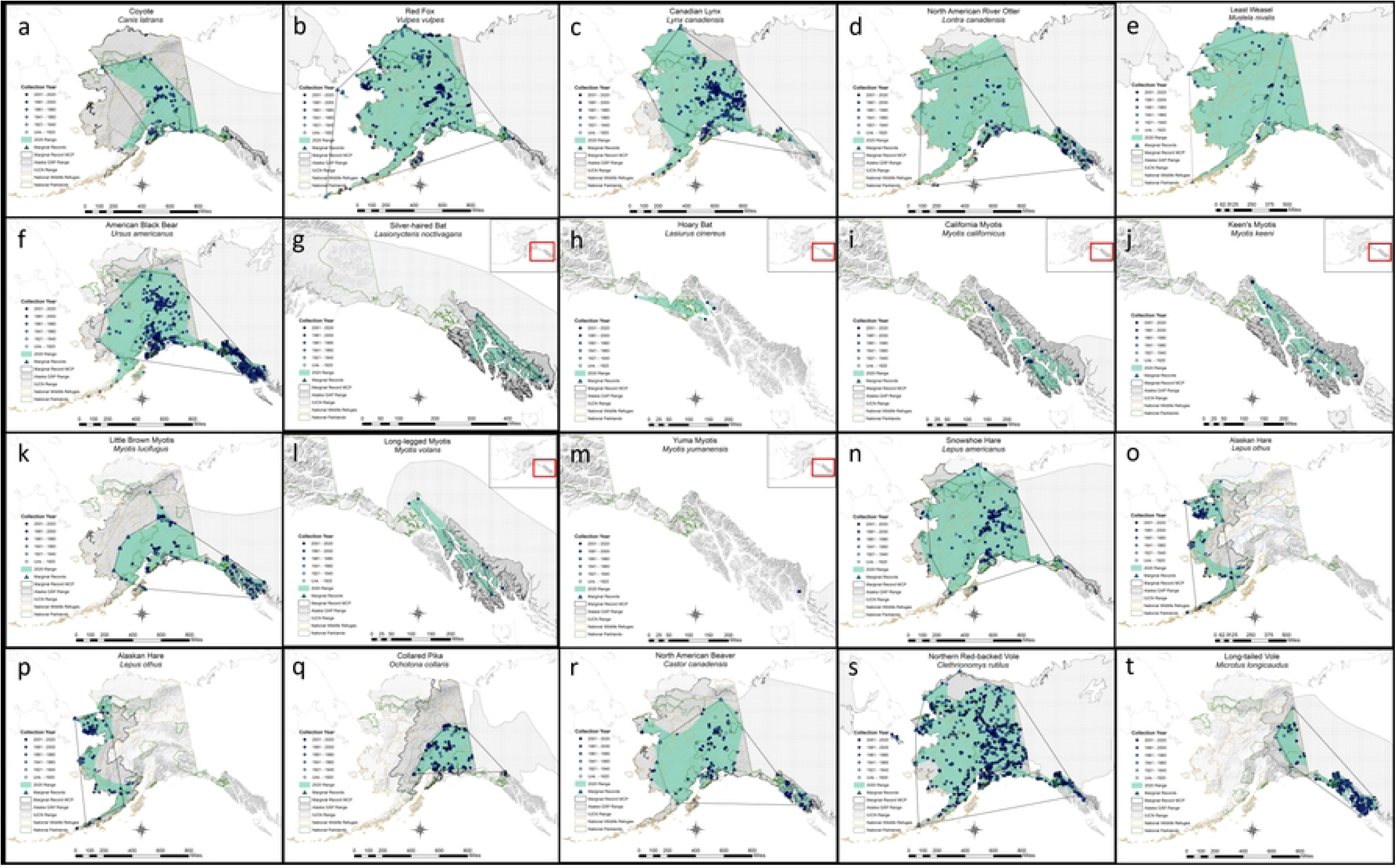
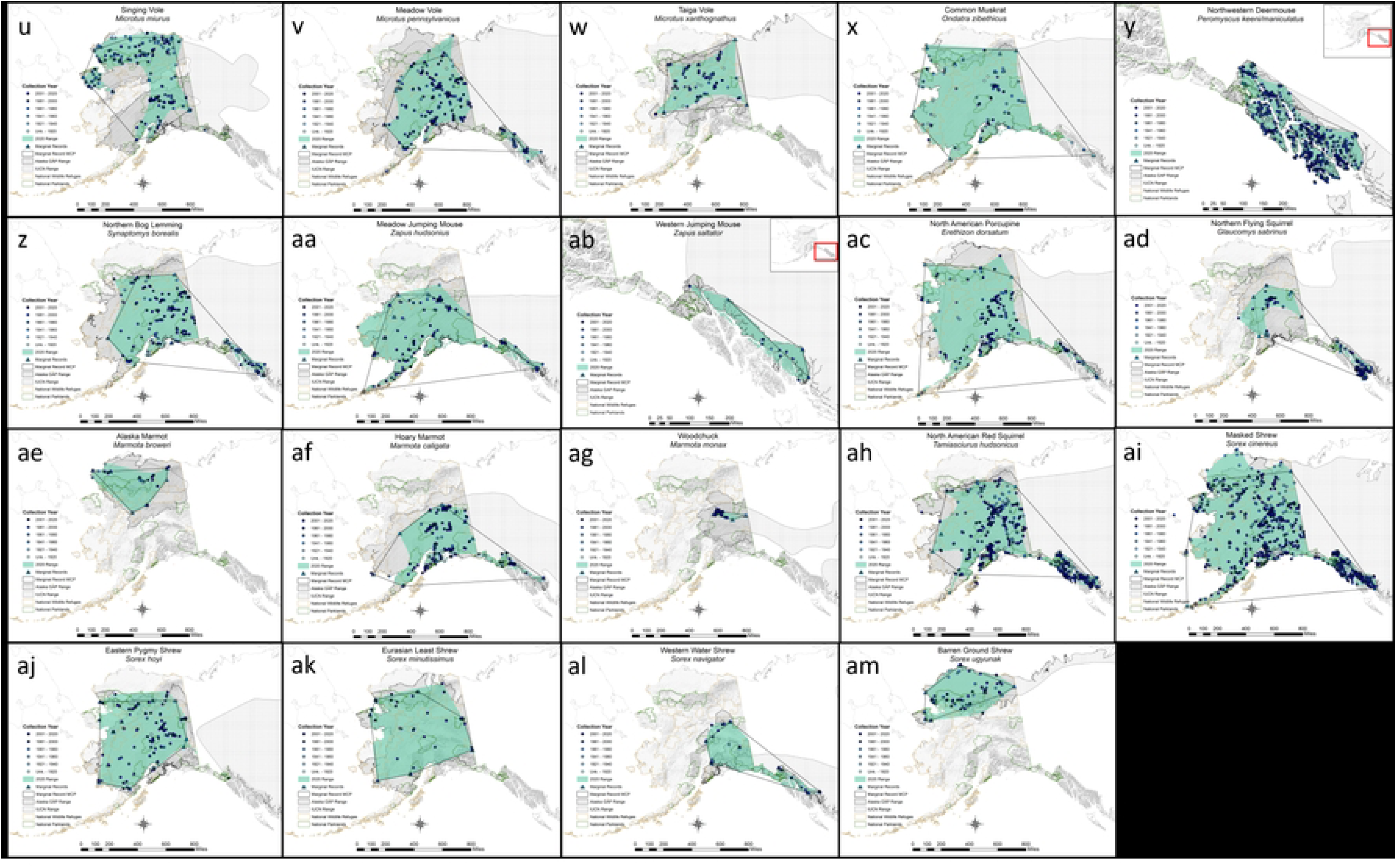
Maps of mammalian occurrence records. Maps for 39 species of terrestrial Alaskan species with extralimital records (those occurring > 5 km beyond marginal range polygons) harvested from the Global Biodiversity Information Facility (GBIF) [34] are included here (a-am) with range polygons from the International Union for the Conservation of Nature (IUCN) [32], the Alaska Gap Analysis Project (AKGAP) [33], and those based on the minimum bounding geometry of marginal records from MacDonald and Cook [21]. Occurrence records are colored by year of collection with darker colors indicating more recent records and lighter colors indicating older records. Public lands managed by the U.S. National Park Service (green) and U.S. Fish and Wildlife Service (brown) are also shown. All maps are in the NAD83 Alaska Albers projection and have an extent of -173.304°, 51.139° to -112.673°, 67.397°, except maps of Southeast Alaska that are shown in reference to this extent (see inset).

### Eulipotyphla

A total of 677 shrew records, belonging to six species, were identified as extralimital (occurring outside MCP species ranges), including 141 that were collected in 2009 or later. Recent extralimital specimens of shrews were collected primarily across the northern half of the state, although some extralimital specimens were identified in Southeast and Southwest Alaska before 2009 as well (Table 1, Fig 2, S2 Table). Recent extralimital specimens of Eurasian Least Shrews (*Sorex minutissimus* [E. A. W. Zimmermann, 1780]) and Eastern Pygmy Shrews (*Sorex hoyi* [S. F. Baird, 1857]) were collected in new areas across the Brooks Range (Noatak NP&P, Gates of the Arctic NP&P, and Arctic NWR [38], exceeding their MCP ranges to the north by ∼245 km and ∼95 km, respectively. Barren Ground Shrews (*Sorex ugyunak* [R. M. Anderson & Rand, 1945]) were collected for the first time in Kobuk Valley NP and near Kanuti NWR over 130 km to the south of the MCP and were the only Arctic species in our analysis with recent occurrences along their southern range margin.

Recent extralimital specimens of other shrews were collected primarily in the vicinity of older records and so were not indicative of range changes. Extralimital specimens of Masked Shrews (*Sorex cinereus* [Kerr, 1792]) were collected in extreme Southeast Alaska, in Southwest Alaska as far as Unimak Island (Alaska Maritime NWR), on Nunivak Island (Yukon Delta NWR), and north of the Brooks Range (incl. Arctic NWR; Table 2, Fig 2). The addition of older extralimital specimens extended knowledge of their range to include the Beaufort Sea coast, as much as ∼310 km to the north. Extralimital specimens of Tundra Shrews (*S. tundrensis* Merriam, 1900) were collected from the Seward Peninsula, in or near the Yukon Delta NWR, Alaska Peninsula NWR, Yukon-Charley Rivers National Preserve, and Arctic NWR, but also occurred near older specimens (Table 2, Fig 2). Western Water Shrews (*Sorex navigator* [S. F. Baird, 1858]) collected in the White Mountains since 2006 added three extralimital specimens and increased their current range as much as 95 km to the northwest (Fig 2).

### Rodentia

We identified 5,446 extralimital records (including 1,312 after 2009) belonging to 17 rodent species that were distributed across the state (Tables 1-2, Fig 2, S2 Table). Extralimital specimens for rodents indicated modest range extensions in several directions (Fig 2), but different patterns emerged for two groups of species, one with range extensions to the west and northwest and a second with extralimital records primarily to the north and northeast.

Several species—North American Beaver (*Castor canadensis* [Kuhl, 1820]), Northern Red-backed Voles (*Clethrionomys rutilus* [Pallas, 1779]), Northern Flying Squirrels (*Glaucomys sabrinus* [G. Shaw, 1801]), North American Red Squirrels (*Tamiasciurus hudsonicus* [Erxleben, 1777]), Long-tailed Voles, Northern Bog Lemmings (*Synaptomys borealis* [J. Richardson, 1828]), and Common Muskrats (*Ondatra zibethicus* [Linnaeus, 1766])—had new extralimital records that pushed inferred range boundaries to the northwest by ∼5-160 km (Fig 2). Extralimital records of North American Beaver occurred along the northern and western extents of the range (Fig 2). Several records between 2004 and 2020 also place beavers in Bering Land Bridge National Preserve and the Yukon Delta NWR, near their western range extent (Table 2). Northern Flying Squirrels, commonly associated with eastern Alaska, were collected as far west as McGrath, AK in 2013 (<u>UAM:Mamm:117637;</u> see [21] for abbreviations used for museum collections) and west of Denali NP&P between 2014 and 2016, extending the known range of this boreal species ∼120 km westward (Fig 2, S2 Table). A North American Red Squirrel (<u>UAMObs:Mamm:254</u>) was photographed on the western Seward Peninsula in 2019, marking the westernmost extent of this species, normally associated with the boreal forest (S2 Table). However, our current range model still regarded this location as an outlier and did not include it within the range (Fig 2). Two Northern Bog Lemming specimens (<u>MSB:Mamm:291027</u>, <u>291034</u>) collected from Kobuk Valley NP in 2014 represent the northwesternmost detections of this species and extend the current known range extent by ∼160 km (Fig 2, S2 Table).

Extralimital records for a second group of rodents, extended inferred ranges primarily to the north: North American Porcupine (*Erethizon dorsatum* [Linnaeus, 1758]), Singing Voles, Hoary Marmots (*Marmota caligata* [Eschscholtz, 1829]), and Eastern Meadow Voles (*Microtus pennsylvanicus* [Ord, 1815]; Fig 2). Recent extralimital records of North American Porcupines extend the current known range ∼200 km north of the MCP, across the Brooks Range, although still within the IUCN and AKGAP ranges (Fig 2) [32,33]. Thirty singing vole specimens collected from two locations near the Beaufort Sea coast since 2009 mark the northernmost records of this species, ∼52 km north of their MCP range (Fig 2). Extensive sampling between 2010 and 2018 yielded 18 hoary marmot specimens from the Yukon-Tanana River Uplands and represent a ∼100 km extension of their current range to the northeast (Fig 2). Similar collection efforts resulted in the detection of an Alaska Marmot (*Marmota broweri* [E. R. Hall & Gilmore, 1934]; <u>UMObs:Mamm:146</u>) in the Arctic NWR in 2011, extending the known range by ∼30 km to the northeast (Fig 2, S2 Table). <u>NEON</u> (National Ecological Observation Network) surveys also detected the northernmost occurrences of meadow voles at Toolik Field Station and in Arctic NWR in 2019, a northerly extension of their current range by ∼105 km (Fig 2).

For some interior species with smaller ranges, recent extralimital records resulted in only small (< 10 km) extensions beyond the MCP range: Woodchucks (*Marmota monax* [Linnaeus, 1758]), Taiga Voles (*Microtus xanthognathus* [Leach, 1815]), and Northern Meadow Jumping Mice (*Zapus hudsonius* [E. A. W. Zimmermann, 1780]; Fig 2). Extralimital records in Southeast Alaska added new islands and areas of the mainland to the inferred ranges of rodent species whose extents terminate in this region, including Northwestern Jumping Mice (*Zapus saltator* [J. A. Allen, 1899]; Fig 2). While we identified 1,047 extralimital records of Northwestern Deermice, analysis for this species could not be separated from that of the closely related Eastern Deermouse (*P. maniculatus*) given their fluid taxonomy. Genetic barcoding is necessary to accurately discern these species (Boria & Blois 2023) and we had low confidence that GBIF records of *P. maniculatus* within the range of *P. keeni* were correctly differentiated.

### Lagomorpha

We report 133 extralimital records for the three species of native lagomorph that occur in Alaska, including 55 since 2009 (Table 1). Extralimital records occurred primarily around the periphery of the state, west- and northward (Fig 2). Older extralimital specimens of snowshoe hares (*Lepus americanus*) were collected across the North Slope (of the Brooks Range), extending their inferred range at least 75 km to the west. Several recent records were identified from the Seward Peninsula, Southwest Alaska, Kodiak NWR, and northern Southeast Alaska (Tables 1-2, Fig 2). Numerous recent and older extralimital records of Alaskan hares (*L. othus*) occurred in Northwest Alaska, including in Selawik NWR and Noatak National Preserve; Cason et al. 2016), resulting in a new range extent ∼185 km to the northeast (Table 2, Fig 2). Older extralimital records originated from the base of the Alaska Peninsula (Katmai NP&P and Alaska Peninsula NWR), whereas newer records came from as far west as Unimak Island (Alaska Maritime NWR; Table 2). The Collared Pika (*Ochotona collaris* [E. W. Nelson, 1893]), whose range is limited to the southeastern quadrant of mainland Alaska, was represented by several recent extralimital specimens from the White Mountains, as much as ∼45 km northwest of their MCP range, and one new observation (<u>iNaturalist:7671018</u>) from near Klondike Gold Rush NHP in 2017. Five specimens collected from Lake Clark NP&P in 2007 mark the westernmost known extent of this species in Alaska (Fig 2, S2 Table).

### Chiroptera

Following the corrected identification of 15 Yuma Myotis (*Myotis yumanensis* [H. Allen, 1864]) specimens collected from southern Southeast Alaska (Olson et al. 2014) and the auditory detection of Northern Hoary Bats (*Lasiurus cinereus* [Palisot de Beauvois, 1796]) at four locations in northern Southeast Alaska in 2012-13, seven species of bat are now known to occur in Alaska (Table 1, Fig 2). In total, we identified 159 extralimital records of bats (118 since 2009; Table 1). With the exception of the Little Brown Myotis (*Myotis lucifugus* [Le Conte, 1831]), all of these were from Southeast Alaska. Recent extralimital specimens of California Myotis (*M. californicus* [Audubon & Bachman, 1842]) and Long-legged Myotis (*M. volans* [H. Allen, 1866]) were collected from new islands in the Southeast Alaska, while extending the inferred northern and southern range extents for both species (Fig 2). The current observed western range limit of Silver-haired Bats (*Lasionycteris noctivagans* [Le Conte, 1831]) now includes the island of Sitka after the photographic observation (<u>iNaturalist:54677027</u>) of an individual there in 2020 (Fig 2, S2 Table). Little Brown Myotis is the only bat known to occur across Interior Alaska (between the Brooks and Alaska Ranges) and the Kodiak NWR [21,39–41]. The northernmost extralimital specimen (<u>UAM:Mamm:127701</u>) was collected in Wiseman, AK (adjacent to Gates of the Arctic NP&P) in 2013, pushing the extent of its known range ∼170 km to the north compared to the marginal MCP (Table 2, Fig 2, S2 Table).

### Carnivora

We identified 436 records (74 since 2009) from six carnivore species as having extralimital records (Table 1, Fig 2, S2 Table). American Black Bears (*Ursus americanus* Pallas, 1780), Red Foxes (*Vulpes vulpes* [Linnaeus, 1758]), and North American River Otters (*Lontra canadensis* [von Schreber, 1776]) had a mixture of recent and older extralimital records in both Southeast and northern Alaska (Table 1, Fig 2, S2 Table). Canadian Lynx (*Lynx canadensis* [Kerr, 1792]) and Red Fox were recently documented in the Alaska Peninsula NWR and Alaska Maritime NWR, respectively, demarking their southwestern range extents (Table 1, Fig 2). Three records of North American River Otters since 2014 in the Brooks Range and on the Beaufort Sea coast in the Arctic NWR are the northernmost records of this species in Alaska. Numerous extralimital records of Red Fox along the northern coast, especially 31 recent occurrences in northeastern Alaska near oil infrastructure and in the Arctic NWR, are representative of a range that extends 200 km to the northeast and now covers nearly all of Alaska. Numerous older extralimital records of Coyotes (*Canis latrans* [Say, 1823]) were identified from the Kenai NWR in the south but also recent records from Gates of the Arctic NP&P and Glacier Bay NP&P regions are indicative of (but likely still underrepresent) a Coyote range that has been expanding in Alaska for at least the past 70 years [21,42]. Least Weasels (*Mustela nivalis* [Linnaeus, 1766]) occur across most of Western and northern Alaska, but a string of older specimens (including in Yukon Flats NWR) and new specimens in Eastern Alaska near Yukon-Charley Rivers National Preserve and Glacier Bay NP&P represent an eastward extension (∼830 km) of the current inferred range of this species.

## DISCUSSION

### Range Extensions

We identified extensions of previously inferred range limits in multiple directions for 39 terrestrial Alaskan mammals including two species (Northern Hoary Bats and Yuma Myotis) that were detected in Alaska for the first time during the past decade (Tables 1-2, S2). While new extralimital records may indicate the actual expansion of a range, they could instead reflect previously unsampled localities where a particular taxon may have long occurred; this can be a decidedly difficult distinction, especially given the recent digitization of older records [43]. This latter consideration best fits many of the historic specimen records not included in MacDonald and Cook’s [21] review. During their assessment, many North American museums had either not yet digitized their collections or were not yet serving data to public online repositories.

Even after accounting for historically sampled yet previously unknown localities, anthropogenic climate change appears to be shifting the distribution of many species [8,44,45]. Our primary aims here were to highlight extralimital records indicative of possible range extensions, to provide an update of the current geographic status of terrestrial mammals on NPS and FWS lands in Alaska, and to offer new, data-driven range maps that represent the current state of knowledge for terrestrial species at the forefront of climate change. While deciphering potential causes for any range shifts is complex and beyond our scope, we can compare observed extralimital occurrences against predictive geographic species distribution models (S1 Table) [22,25] and statistical habitat models [4] that predict where and how distributions of mammals were likely to change over the coming century. Below, we consider recent (i.e., after 2009) extralimital records in the context of available predictions of small mammal species for the 21^st^ century [4,22,25].

### Eulipotyphla

Recent extralimital records of shrews were identified along Alaska’s west coast and across the Brooks Range (Fig 2). Most extralimital specimens were collected in the vicinity of older records and likely represent more complete occurrence datasets rather than shifts in species ranges since 2009. However, specimens of Least, Pygmy, and Barren Ground Shrews were identified to the north and east of extents inferred from MacDonald and Cook’s [21] maps. Hope et al.’s [25] current models generally predicted near statewide (except the eastern Brooks Range and Northeast Alaska) distributions for Masked, Pygmy, Tundra, Least, and Western Water Shrews. Although models for these species were largely under-predictive for the current (2015) timeframe, model trends indicated expanding distributions for most shrews into at least the southern Brooks Range during the 2020s. Baltensperger and Huettmann’s [22] models also predicted range expansions for these species onto the North Slope but contractions in Southwest Alaska by 2100. In contrast to Hope et al.’s [25] models, Baltensperger and Huettmann [22] indicate a small present-day (2015) range for Western Water Shrews that was predicted to expand northwestward across the state by 2100. Recent extralimital specimens for Western Water Shrews are consistent with this prediction. Both sets of models were in general agreement for most species, except Barren Ground Shrews. Baltensperger and Huettmann [22] predicted the range of this species to recede north of the Brooks Range by 2100, whereas Hope et al.’s [25] models predicted a gradual southward expansion of the range by the 2080s, a prediction consistent with the most recent extralimital record (<u>MSB:Mamm:143102</u>) for this species along the southern margins of the Brooks Range.

### Rodentia

Observed extensions of previously inferred range margins for the northwestern group of rodents are consistent with future distribution models that predicted northwestwardly expanding distributions of Long-tailed Voles, Northern Bog Lemmings, Northern Red-backed Voles [22] and North American Red Squirrels [25]. Statistical projections also predicted increased areas of available habitat for Northern Flying Squirrels, North American Red Squirrels, and Northern Red-backed Voles, whereas habitat areas for most other species were predicted to remain roughly the same [4]. Hope et al.’s [25] models predicted North American Red Squirrels to occur on the Seward Peninsula by the 2050s, where an individual was recently observed near Nome, AK (although no other records have since been reported). Alaska Marmots were also recently detected in an area predicted by the 2080s to be the last refugium for this species [25]. Despite a predicted decline in habitat available for North American Beavers [4], several recent extralimital records and a comprehensive analysis of historical observations indicated that the species has, in fact, been expanding its distribution across northern and western Alaska [31]. New extralimital specimens of Northern Bog Lemmings in Kobuk Valley NP were collected from areas where this species was predicted to occur over the coming decades [22,25].

Results for the second group of rodents (those with recent extralimital records primarily to the north and east; i.e., North American Porcupine, Meadow Vole, Singing Vole, Hoary Marmot), were mixed relative to prior model predictions. Newly documented occurrences of North American Porcupines 28−200 km north of their previous MCP range extent are consistent with Marcot et al.’s [4] projection that their habitat would increase over the coming century. Both Marcot et al. [4] and Baltensperger and Huettmann [22] predicted a decline in total available habitat and area of occurrence for Singing Voles and Meadow Voles. Despite overall predicted declines in Singing Vole distribution, Baltensperger and Huettmann’s [22] models indicated a small northward extension of their 2100 distribution that includes the Beaufort Sea coastline, the same region from which several extralimital specimens were recently collected. Recent extralimital records of Meadow Voles ∼25−112 km north of their MCP range lend support to Hope et al.’s [25] model that indicated an expanding distribution. Hope et al. [25] predicted a small northward expansion of Hoary Marmots by the 2080s, but models underrepresented occurrence records and did not include the area of eastern Interior Alaska where new extralimital records were recently documented ∼35-135 km north of the MCP.

### Lagomorpha

Extralimital records of Snowshoe Hares largely conformed to Hope et al.’s [25] distribution, though our current range map indicates broader occupation in Interior Alaska, a modest extension to the west and southeast (Fig 2). These results are consistent with trends of increasing shrub cover in tundra areas that should make these areas more hospitable for Snowshoe Hares [28], although this contradicts future habitat projections that indicated no significant change in available habitat [4]. Recent extralimital records of Alaskan Hares indicated the occurrence of this species along the length of the Alaska Peninsula, and to north of the MCP [27]. In contrast, Hope et al.’s [25] models did not predict the Alaska Peninsula to be part of the Alaskan Hare’s range during the 21st century. New records from the Seward Peninsula are consistent with predictions of an expanding distribution across northern Alaska [27], but there were no recent extralimital records to indicate inland movement of their range as predicted by Hope et al. [25]. In contrast, Marcot et al. [4] predicted a small but insignificant decline in available habitat for Alaskan Hares. Distribution models for Collared Pika predicted a constriction of their range over the coming decades [25], whereas recent extralimital records represent a small extension of their current observed range to the north and south, which may only be a product of expanded sampling (Fig 2).

### Chiroptera

Detection of bat species in Alaska has recently been augmented using acoustic recording to detect species in new locations, including the detection of Northern Hoary Bats, and the collection of Yuma Myotis in Alaska for the first time during the past decade. While new detections have expanded occurrence sets of bats in southeast Alaska [46,47], modeled trends in distribution, range, or available habitat at regional to statewide scales have not yet been conducted for mainland Alaska (but see [48,49]). This makes evaluating whether new extralimital records represent actual range expansions or just improved sampling an even greater challenge. Given the small size of bat occurrence datasets, the difficulty in detecting bats in the field in many Alaskan habitats, and the technological developments that have made this work more feasible only recently, we suspect that new records of Chiroptera (more than other orders) represent a growth in the known occurrence set, rather than movements of species. More detection and modeling research is needed to improve our collective conception of the geographic status of bats in Alaska and we conjecture that further auditory detection efforts will continue to identify bat records in many new locations, especially on islands in Southeast Alaska.

### Carnivora

Among the six species of carnivore with extralimital records (Red Fox, Coyotes, Canadian Lynx, American Black Bears, North American River Otters, and Least Weasels), all exhibited either a westward and/or northward extension of range boundaries, whereas Canadian Lynx and Red Fox were the only species without extralimital records to the southeast. According to projections by Marcot et al. [4], the area of suitable habitat was predicted to grow significantly for American Black Bears (> 10 %), stay about the same for Red Fox, Coyotes, and Canadian Lynx, but decline (> 10 %) for North American River Otters (Least Weasels were not modeled). Habitat predictions were largely consistent with the extralimital datasets reported here, although new records of North American River Otters indicated a sizeable extension of their range that now includes most of the North Slope and new portions of Southeast Alaska, contradicting projected habitat trends [4]. Spatially explicit models predicting distribution change among members of Carnivora were conspicuously lacking and we encourage research in this area to contextualize new and recently documented extralimital records.

### Extralimital Records on Federal Lands

Among national parklands in Alaska, we identified eight NPS units—Lake Clark NP&P, Glacier Bay NP&P, Klondike Gold Rush NHP, Bering Land Bridge National Preserve, Gates of the Arctic NP&P, Kobuk Valley NP, Noatak National Preserve, and Cape Krusenstern NM—as containing recent extralimital records (≥ 2009) for 11 mammal species (Coyote, American Beaver, North American River Otter, Keen’s Myotis, Masked Shrew, Eastern Pygmy Shrew,Eurasian Least Shrew, Barren Ground Shrew, North American Red Squirrel, Northern Bog Lemming, and American Black Bear; Table 2, Fig 2). While these records may not represent new species for NPS units, consistent monitoring programs could be used to provide indications of species that may be expanding in these areas.

Eight National Wildlife Refuges—Alaska Maritime NWR, Alaska Peninsula NWR, Becharof NWR, Kodiak NWR, Togiak NWR, Yukon Delta NWR, Selawik NWR, and Arctic NWR—contained recent extralimital records for 17 species (Beaver, North American Porcupine, Snowshoe Hare, Alaskan Hare, North American River Otter, Canadian Lynx, Hoary Marmot, Alaska Marmot, Singing Vole, Meadow Vole, Masked Shrew, Eastern Pygmy Shrew, Eurasian Least Shrew, Tundra Shrew, Barren Ground Shrew, North American Red Squirrel, Red Fox; Table 2, Fig 2). All units are located around the periphery of the state, in the West and North, where most terrestrial mammals in our analysis had extralimital records. The Arctic NWR contained by far the most species (n = 13) with recent extralimital records, indicating the increasing importance of this region for its growing biodiversity and as a destination for northward-moving boreal species.

### Inferring Range Shifts from Occurrence Data

Museum specimen and citizen science data enhance spatial analyses, biological monitoring, and conservation assessments [50]. Both range maps and distribution models, as well as subsequent geospatial and biodiversity analyses, depend on accurate, verifiable georeferenced occurrence datasets as their basis. Ultimately, the quality, completeness, and availability of these records dictate the accuracy, relevance, and timeliness of analyses and management decisions. While extralimital records identified here may seem to generally conform to prior model predictions for small mammals, is the accuracy and completeness of the terrestrial mammal record set sufficient for evaluating changing range extents? How confident can we be that any “extralimital” record does, indeed, represent an actual range extension?

Records harvested from GBIF generally corresponded well with MacDonald & Cook’s [21] assessments, but new and newly digitized observations indicated possible range extensions > 5 km for 39 species. Over 1,700 recent extralimital records hint at the possibility that the range extensions summarized here may represent real movements and are not only artifacts of sampling effort or delayed digitization. Among initially identified extralimital records, just 4% contained errors and were flagged as “dubious.” However, we only checked records along range margins for accuracy, and numerous georeferencing and taxonomic identification errors likely remain in archives and online repositories. Additionally, given the lag between specimen collection, expert identification, and data digitization, new species occurrences or specimen-based range extensions resulting from recent field work and literature may not yet be reflected in recent GBIF data harvests. Finally, while museum and citizen science records provide a baseline for species presence, they are inapplicable to the documentation of species absences. Establishing better mechanisms for documenting and sharing absences of expected species would provide a better baseline for shifting distributional data. Only increased sampling and specimen archival from understudied regions of the state (e.g., Southwest and Western Alaska; Fig 1) will help to improve confidence that we are working with comprehensive and representative geospatial datasets, yet the difficult logistics of conducting field work in remote Alaska underlie slow progress towards this goal.

To differentiate anecdotal observations from verifiable records, some type of physical record allowing for the positive identification of a species must be made available. We echo prior calls to explicitly recognize and rank evidentiary standards and advocate for the highest level of reliable evidence for the rarest species, while allowing other types of verifiable documentation for more common species [51]. The most verifiable practice for identifying and archiving mammals is the collection and preservation of whole specimens, including skulls (dentition is necessary for differentiating many small mammals). This is particularly important for assessing species that have undergone taxonomic revisions or are difficult to identify, e.g., *Martes americana/caurina* (Pacific marten [Merriam, 1890]), *Peromyscus keeni/maniculatus*. With the rise of digital citizen-science repositories, a sufficient level of verifiability may be met with a physical specimen, DNA or eDNA sample, or quality photograph (although most North American small mammals cannot be reliably identified to species based on photos) [52]. For example, a photo that has been identified and confirmed by at least 2/3 of reviewers is necessary for iNaturalist records to become “research grade” and meet the standards for GBIF archival. In some cases, diagnostic auditory evidence (e.g., bats, canids, pikas, marmots, etc.) and photographs or impressions of tracks (e.g., carnivores, ungulates) are also sufficient and useful for the identification of some common or readily diagnosed species that may be difficult to observe otherwise. While some museums serve these types of observational data from their databases, community standards for distribution and access to disparate observational data types are still being developed.

We also advocate for adherence to the FAIR principles of scientific data management and stewardship: Findability, Accessibility, Interoperability, and Reuse of digital assets [53]. To achieve a sustainable level of completeness for the global occurrence datasets of species, all natural history records should ideally be georeferenced, digitized, and served to a central, public repository [54]. Currently, over 1,600 institutions contribute records to one of these online repositories, but numerous federal and private institutions do not, at least on a consistent basis. To this end, iDigBio and BISON have been instrumental in digitizing older collections. Streamlining the dataflow from field observations to accurate, complete, and accessible digital collections is a goal that field biologists, management agencies, and natural history repositories alike can and should strive towards. Through this common workflow, we may improve confidence in the geospatial datasets that form the basis of range maps, distribution models, and quantitative analyses that inform management recommendations and conservation priorities.

### Improving Range Map Accuracy

Freely accessible range maps are a critical tool for assessing biodiversity patterns [55], but considerable gaps in coverage and questions about accuracy of specific maps remain [37]. For example, most range maps did not accurately reflect the extent of geographic records for 6 species (Keen’s Myotis, Long-legged Myotis, Yuma Myotis, Long-tailed Vole, Northern Flying Squirrel, and Hoary Marmot; Table 1, Fig 2). Further research into the recent biogeographic history of these species as well as new field-detection efforts are needed to further clarify and track changes. Ultimately, many of the reviewed range maps are served to inform the public about species biogeography and they are also used extensively by scientists who include them in scientific presentations, incorporate them into spatial analyses, and ultimately to inform conservation assessments (e.g., [56]). As such, current and accurate data underpinning these maps is paramount.

Accurate range maps should attempt to balance errors of omission and commission. The AKGAP range maps [33] encompassed most species occurrences, but at the 8-digit HUC watershed scale tended to overestimate occupancy. Regional GAP projects across the country have been valuable for identifying biodiversity hotspots and potential connectivity between disjunct distributions (e.g., [57]). While sensitivity of AKGAP range maps to occurrence data was generally quite good, specificity was poor (< 50.0%) for 13 species: Coyote, North American River Otter, Least Weasel, Little Brown Myotis, Collared Pika, North American Beaver, Singing Vole, Common Muskrat, North American Porcupine, Northern Flying Squirrel, Hoary Marmot, and Eurasian Least Shrew (Table 1). Such mismatches between occurrences and watershed occupancy are the result of either a lack of records in under-sampled areas or range polygons that overestimated the geographic extent of these species [33]. As such, we support the use of smaller watershed units (e.g., HUC12) to delineate range extents (e.g., [58]). In contrast, IUCN range maps encompassed fewer than 90.0% of the occurrence records at the statewide scale for more than half of analyzed species with extralimital records (Table 1). Methods guiding the development of IUCN range maps encourage species assessors to provide the best possible map based on available data, including occurrence point data, polygon data, and watersheds [32]. However, expert construction of range maps for many species did not appear to be well matched to the occurrence data in Alaska. Ensuring range maps used in spatial analyses, conservation assessments, and management decisions are accurate at statewide-to-regional scales has important relevance for the many downstream uses of these data.

Ideally, range maps will be updated in real or near-real time as new records are reported, so that archived maps may serve as a dynamic and reliable historic record of biogeographic change. This can be accomplished by improving species mapping and assessment processes so that species are regularly reviewed in a well-documented, referenced, and repeatable manner, which is key to providing better baseline biodiversity data. While having maps created or vetted by taxonomic experts can improve map quality [59], this is not always possible for all species. The option we implemented here was to analyze available occurrence records with a common range mapping tool with a limited number of settings [35]. Consistent use of such a tool to map the concave range extents of multiple species while stipulating how parameters are set will help to remove subjectivity and overgeneralization from expert range maps and go a long way towards standardizing range maps across species and time.

## CONCLUSIONS

The actual and perceived ranges of species are continually changing, both as we assimilate occurrence records into increasingly networked and comprehensive data repositories and as species move in response to climate change and other anthropogenic and competitive stressors. Extralimital records and newly developed minimum concave range maps highlighted inferred range extensions as compared to MCP range polygons for 39 terrestrial mammal species in Alaska. As a disproportionate number of extralimital records were also identified from lands managed by the U.S. National Park Service and U.S. Fish & Wildlife Service in western and northern Alaska (especially Arctic NWR), these areas may serve as an early indication of changing mammalian faunal systems in Alaska as well as provide clear opportunities for monitoring species shifts. While it remains difficult to identify precise causes for changing species geographies, new extralimital observations were largely consistent with prior model predictions of northward and coastward movement of many boreal species over the coming century. To monitor changes in the geographic arrangement of species over time, it is imperative to continue field sampling efforts along range margins and to contribute voucher specimens and accurate occurrence data to appropriate public repositories so that we can use consistent range mapping methods (as proposed here) to regularly update species range maps and models over the coming century.

## ACKNOWLEDGMENTS

We heartily thank all the natural history museums and their curators, collection managers, and collectors, especially Bob Timm (University of Kansas Natural History Museum), Chris Conroy (UC Berkeley Museum of Vertebrate Zoology), Gary Shugart (Puget Sound Museum of Natural History), Jeff Bradley (Burke Museum), John Demboski (Denver Museum of Nature & Science), Jon Dunnum (Museum of Southwestern Biology), Kamal Khidas (Canadian Museum of Nature), Shannen Robson (Natural History Museum of Los Angeles County), Kristof Zyskowski (Yale Peabody Museum), Mark Omura (Museum of Comparative Zoology, Harvard University), Maureen Flannery (California Academy of Sciences), Dawn Roberts (Chicago Academy of Sciences), Elmer Finck (Sternberg Museum of Natural History), Melanie Bucci (University of Arizona Museum of Natural History), Eric Rickart (Natural History Museum of Utah), Cody Thompson (University of Michigan Museum of Zoology), and Aren Gunderson (University of Alaska Museum), and all the citizen scientists who contributed to this work in Alaska. We are particularly grateful to Stephen O. MacDonald and Joseph A. Cook, whose landmark 2009 book inspired this study, to Ken D. Tape for comments on a previous version of this manuscript, and to Marissa Breslin for formatting and editing assistance.

## COMPETING INTERESTS

The authors declare there are no competing interests.

## AUTHORSHIP

L.E.O. and A.P.B. conceived the idea for this review; A.P.B. analyzed the data, created the maps, and led the writing; L.E.O. reviewed the maps and contributed to the writing. H.C.L contributed to the writing and conceptual development improvement.

## FUNDING

This work was funded by a University of Alaska Centennial Postdoctoral Award to A.P.B. and generous support from the Jay Pritzker Foundation to L.E.O.

## DATA AVAILABILITY STATEMENT

All maps, including occurrence record locations and range polygons as well as a list of extralimital records and attributes are published online in the Open Science Framework at: https://osf.io/h6u5n

## SUPPLEMENTARY FILES

S1 Table. **Overview of the changing nomenclature of terrestrial Alaskan mammals.** Scientific and common names of species reviewed in this research. Columns indicate whether each species (and its nomenclature at the time) was reviewed by MacDonald & Cook [21] Alaska Gap Analysis Project (AKGAP) [4,22,25,33]. IUCN Assessment indicates the most recent year of review for each species [32].

S2 Table. **Extralimital records for 39 species of terrestrial Alaskan mammals.** Collection and archival attributes (with links to museum records) are included for each specimen reviewed in this analysis.

## Notes

### Competing Interest Statement

The authors have declared no competing interest.

